# DISCO-LAMP: A Novel discontinuous LAMP assay for isothermal antigen detection

**DOI:** 10.64898/2026.01.23.701152

**Authors:** Benjamin M Thomas, Rudo A Simeon, Katherine L Yan, Vikas Chonira, Wesly T Chen, Emma L Webb, Caitlyn Mutchler, Alexi M Fernandez, Jinu Han, Zhilei Chen

## Abstract

Proximity ligation assay (PLA), in which the ligation of two DNA probes is greatly accelerated by the associating target molecules, has emerged as a highly sensitive technique for protein detection. The detection of the ligated DNA typically relies on PCR, which requires temperature cycling. In this study, we report on a novel <u>disco</u>ntinuous (DISCO)-LAMP assay that enables the wash-free detection of PLA products via loop-mediated isothermal amplification (LAMP). Due to the exponential amplification nature of LAMP, a careful balance between efficient amplification of the ligated full-length DNA and minimal background amplification from the individual constituent probes is essential but often challenging to achieve. After extensive template/primer design and assay optimization, DISCO-LAMP assay achieved a detection limit of 1 fM for the ligated DNA probe while maintaining undetectable background amplification at 1 nM of each individual probe. DISCO-LAMP detected Shiga toxin 2 (Stx2) with a limit of detection (LoD) of 100 fM when functionalized with Stx2-binders, as well as both Wuhan-1 and Omicron spike protein when functionalized with D_S16_, a newly engineered DARPin targeting a conserved epitope on the SARS-CoV-2 Spike protein. We believe DISCO-LAMP represents a versatile and efficient LAMP-based PLA technology that is readily adaptable for sensing diverse targets.

## Introduction

Sensitive, accurate and reliable detection of proteins in solutions, such as toxins, biomarkers and antibodies, is a cornerstone for clinical medicine. In the past decades, proximity ligation assay (PLA) has emerged as a highly sensitive and specific analysis method for detecting protein-protein interactions, post-translational modifications, as well as individual proteins directly^1^. In PLA, the coordinated binding of a target molecule by a pair of DNA probes leads to a substantial increase in the local concentration of these probes—by approximately 10^4^-fold—which promotes the ligation of the DNA probes, enabling the detection of the target molecule through amplification of the ligated DNA probes. PLA provides a highly sensitive and specific means for the detection of pathogens^2^, cytokines^3^, cellular proteins/RNAs^4, 5^ and diverse non-nucleic acid molecules^6^.

The most common assay for detecting/quantifying ligated DNA probes is quantitative PCR (qPCR) employing a pair of primers that anneal to each DNA probe^7-9^. Only ligated DNA probes can be efficiently amplified and detected with a pair of primers targeting each probe. However, qPCR requires bulky and expensive thermal cyclers and is not suitable for field applications. To bypass the need for temperature cycling, a number of isothermal DNA amplification methods have been exploited. The most common isothermal DNA amplification method used in conjunction with PLA is rolling circle amplification (RCA) in which PLA mediates the circularization of a DNA molecule which serves as the template for synthesizing long single stranded DNA (ssDNA) molecules. RCA requires a DNA polymerase capable of strand displacement and typically uses Phi29 DNA polymerase derived from *Bacillus subtilis* phage Ф29 with an optimal operating temperature of 37°C^10^. RCA has been extensively used with PLA for detection of cellular proteins^11, 12^, glycoRNA^13^, as well as soluble biomarkers ^14, 15^.

Another commonly used isothermal DNA amplification method is LAMP, short for Loop-mediated isothermal amplification^16, 17^. Both RCA and LAMP use a strand-displacing DNA polymerase. However, unlike RCA, which requires a single primer to linearly amplify the DNA template, LAMP uses 4-6 primers and exponentially amplifies the DNA through the formation of dual-hairpin (dumbbell) structures^16^. Consequently, LAMP can generate up to 10^9^ copies of DNA within 30 minutes, much faster than RCA which typically takes 1-2 hours^10^. Due to its high specificity, efficiency, and the ability to operate at a constant temperature, LAMP has been widely used in diverse fields, particularly for rapid diagnostics and point-of-care testing^18^. A number of LAMP-based assays have recently been approved by the FDA, including an at home colorimetric COVID-19 & Flu Test^19^.

However, despite the potential advantages of LAMP over RCA, there are few reports of PLA-LAMP assays^20, 21^. Because LAMP amplifies DNA exponentially with multiple primers, even minor off-target binding can quickly cause unwanted amplification. This makes it difficult to design PLA-LAMP templates and primers that minimize background amplification before PLA ligation. In this study, we report a DISCONTINUOUS (DISCO)-LAMP design suitable for PLA-LAMP assays. The LAMP template DNA is split unevenly into two fragments and can only be efficiently amplified upon ligation (*i*.*e*. 1 fM). Minimum background signal is observed even when a single probe is present at 10^7^-fold higher concentration (*i*.*e*. 10 nM). Using a pair of previously engineered protein binders to Shiga toxin 2, – DARPin SHT (D_SHT_) and nanobody G1 (N_G1_) – DISCO-LAMP achieved highly sensitive detection of Stx2 with LoD of 100 fM. Additionally, we report the engineering of DARPin S16 (D_S16_) specific for SARS-CoV-2 spike protein and its use for sensitive detection of spike proteins from both Wuhan-1 and Omicron strains. DISCO-LAMP represents a highly versatile platform for protein detection in diverse diagnostic applications.

## Results and Discussion

### Design of LAMP template and primers

A standard LAMP reaction employs two outer primers (*i*.*e*. F3, B3), two inner primers (*i*.*e*. FIP, BIP) and a DNA polymerase with strand-displacing activity (*e*.*g*. Bst 2.0, **Fig 1A**)^33^. BIP harbors two sites – B1c and B2 – which are identical and complementary to B1c and B2c sequences on the template DNA, respectively. The DNA synthesized from primer B3 (orange) displaces BIP DNA product (green) from the template DNA and renders it single-stranded with a hairpin structure at the 5’ end. Annealing of primers FIP and F3 to the BIP product generates a single-stranded FIP DNA (red) with hairpin structures on both termini. This dual-hairpin (dumbbell) DNA serves as the starting point for exponential amplification, yielding a large quantity of concatemer DNA products that can be quantified and visualized by many fluorescent and colorimetric reagents^18^. The addition of loop primers (*i*.*e*. LB and LF), which bind the hairpin region, has been found to further accelerate the LAMP kinetics^22^. To achieve LAMP-PLA, the LAMP template needs to be split into two fragments, which are reconstituted upon ligation.

**Figure 1.**
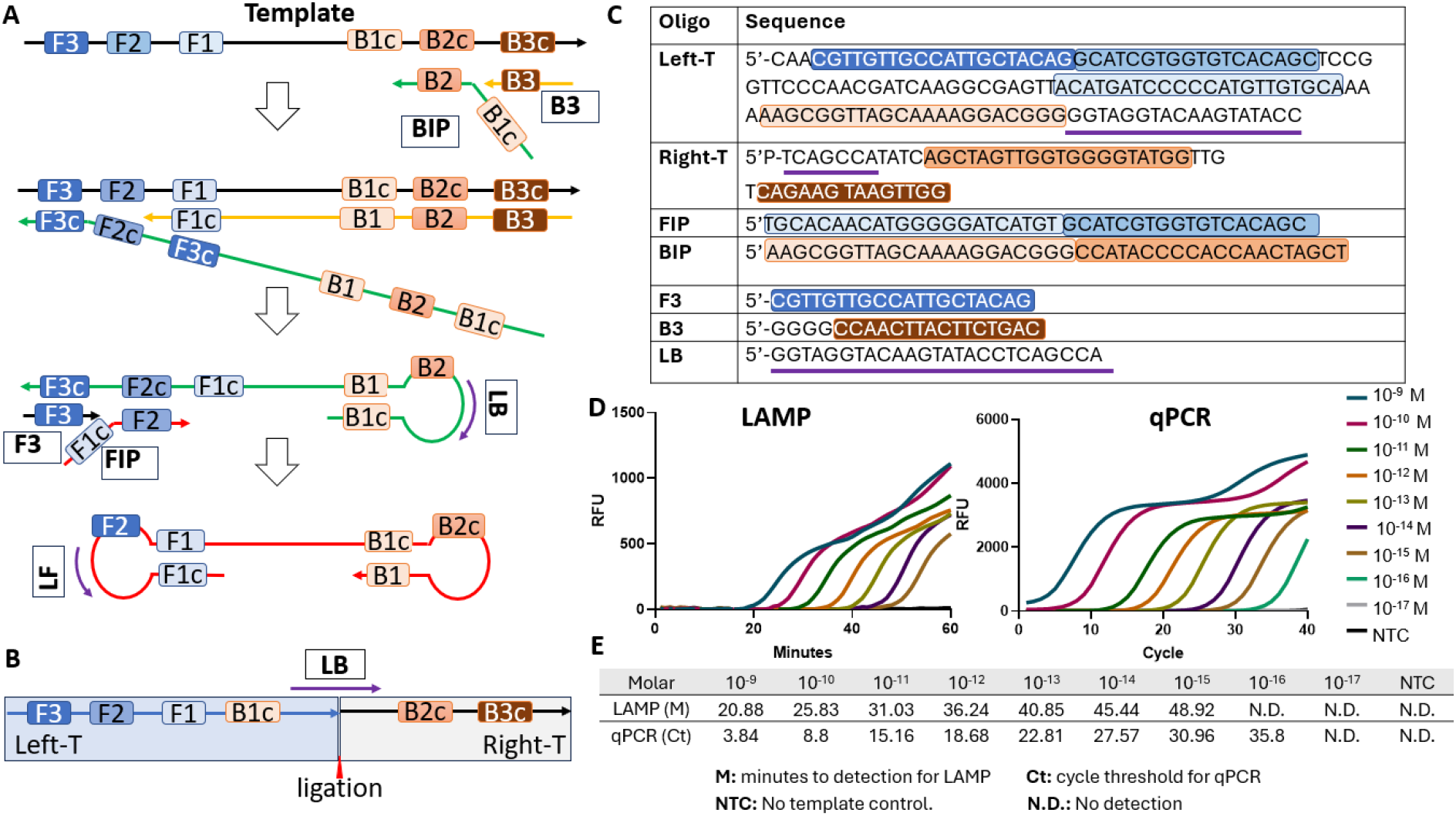
Overview of DISCO-LAMP. (**A)** Overview of LAMP. Primer names are boxed. **(B)** Design of DISCO-LAMP. **(C)** Sequences of DISCO-LAMP template and primers. The annealing regions of different primers are shaded in the respective color. **(D)** Comparison of amplification efficiency of full-length template between qPCR and LAMP. Representative amplification curves from at least 2 experiments were shown. **(E)** Summary of minutes to detection for LAMP (M) and cycle threshold for PCR (Ct).

The design of PCR-PLA template is straightforward, with two primers each annealing to one half of the template. However, LAMP template and primer design need to take into consideration multiple factors such as the primer distance, stability, and potential for primer dimer formation^23^, prompting the development of multiple bioinformatics tools to design LAMP primers (*e*.*g*., PrimerExplorer, NEB LAMP). Unfortunately, none of the existing algorithms consider the possibility of LAMP amplification with half of the DNA template, which we discovered to be a significant challenge due to the nature of exponential amplification. We chose a DNA fragment derived from the β-lactamase resistance gene and our initial design split the template in the middle. However, despite multiple attempts at designing primers to minimize primer dimer formation, as well as tweaking the template DNA and primer sequences to minimize non-specific primer annealing, we were unable to eliminate background LAMP amplification with only half of the template.

Finally, after multiple iterations, we developed a new LAMP-PLA design in which half of the LAMP template – Left-T – harbors regions F3 through B1c, while the other half – Right-T – harbors regions B2c and B3c **(Figure 1B, C)**. Importantly, a loop primer (*i*.*e*. LB) is designed to anneal at the ligation junction. This design enabled the ligated DNA template to be efficiently amplified by our LAMP primer set with a detection limit of 1 fM **(Figure 1D)** and minimum background amplification with up to 10 nM of individual Left- or Right-T DNA probe for 60 minutes with LAMP **(Figure S1)**. No LAMP amplification of the ligated DNA template was observed at concentrations below 1 fM, suggesting a pseudo Yes/No amplification cut-off for LAMP at this concentration. The detection limit of DISCO-LAMP for full-length DNA is slightly worse than conventional LAMP assays and qPCR but nevertheless is highly sensitive.

As a first step to optimize PLA efficiency, Oligos D1, D2, and ND2 are synthesized to model proximity induced by the sensing of a target. D1 and D2 share a 16-nucleotide complementary region while ND2 lacks the complementary region (**Figure 2A, Figure S2A**). Oligo D1 is hybridized to Left-T and initiates the synthesis of double-stranded(ds) DNA with the help of Bst2.0 polymerase. The resulting dsDNA forms the Left-probe. Similarly, Oligo D2/ND2 was hybridized to Right-T to form the Right-probe. The 3’ end of Left-T contains a short sticky end that anneals to D2/ND2 for mediating ligation to Right-T. To evaluate the impact of salt, 100 pM Left-probe (D1) and 10 nM Right-probe (D2 or ND2) were mixed in Buffer L (1x T4 DNA ligase Buffer and 0.1 U/µL T4 DNA ligase) in the absence or presence of 1x PBS. The mixtures were incubated at room temperature for varying times (1-30 minutes), heat inactivated at 70°C for 15 minutes, and analyzed using qPCR and LAMP. With a sticky end of 5 nucleotide, the presence of PBS appeared to significantly improve the PLA efficiency, leading to much larger ΔCt and ΔMinutes between samples with D2 and ND2 regardless of the ligation time (**Figure 2B, C**). Using 1xT4 ligase buffer supplemented with 1xPBS, and 10 minutes ligation time, we also evaluated the impact of ligase concentration and the length of sticky end on the PLA efficiency **(Figure S2B, C)**. We found that the 5-nucleotide sticky end gives the biggest ΔCt between D2 and ND2, although 4- and 6-nucleotide sticky ends also yield strong signal-to-noise ratios. Balancing between PLA efficiency and ease of assay operability/reproducibility, we selected 30 minutes ligation time and 0.1 U/µL T4 ligase as the condition for subsequent studies.

**Figure 2.**
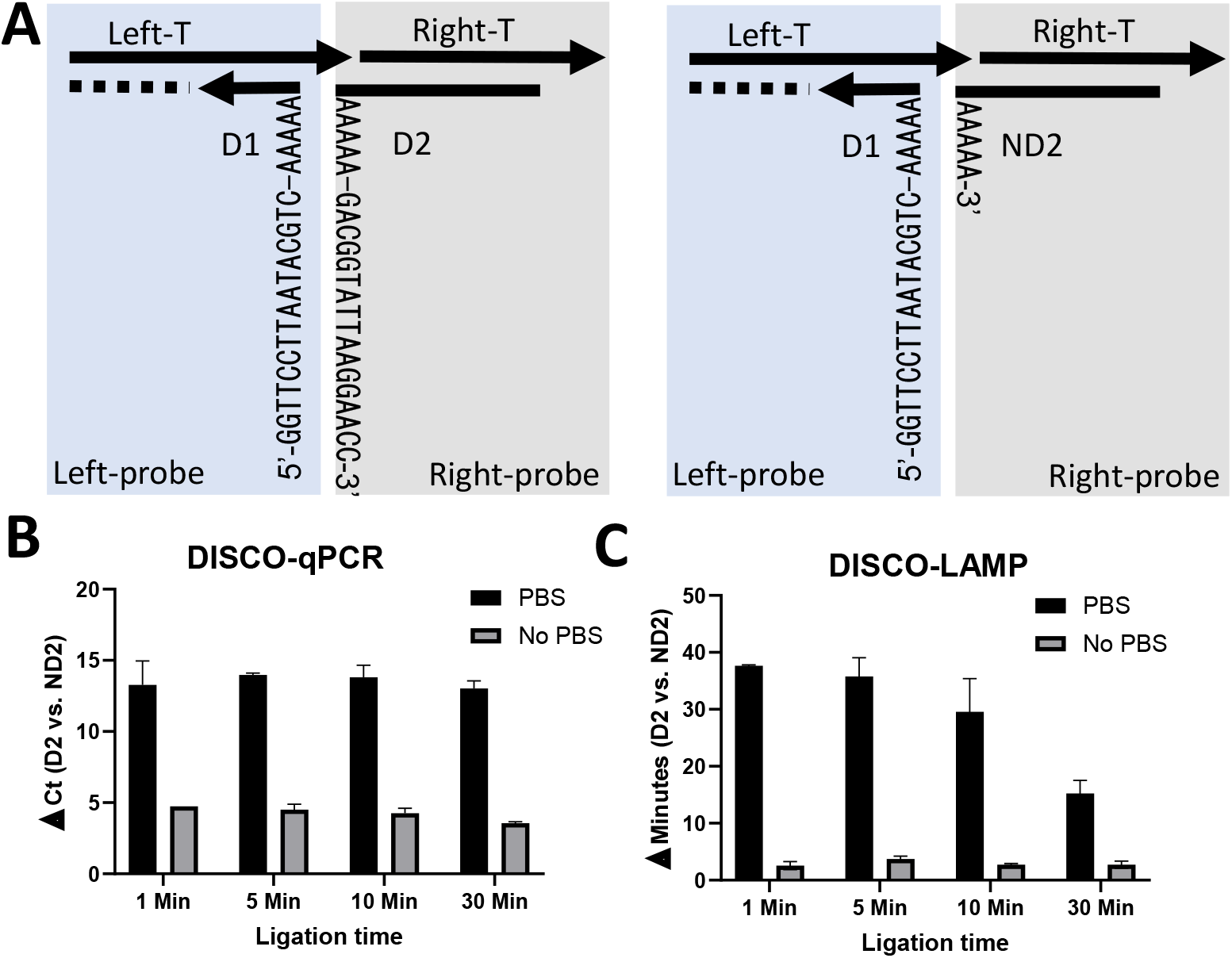
DISCO-LAMP proof of concept. **(A)** Schematic of DISCO-LAMP using oligos to model target sensing. PLA efficiency in 1xT4-ligase buffer alone or supplemented with 1xPBS and analyzed by qPCR **(B)** or LAMP **(C).** Bar graphs show the mean values from two biological replicates, and error bars indicate the standard error of the mean (SEM).

Our DISCO-LAMP target protein sensing assay is analogous to a conventional sandwich ELISA and employs a pair of non-competitive target molecule binders **(Figure 3A)**. For target sensing of proteins, Oligos BL and BR are each synthesized with a dibenzocyclooctyne (DBCO) molecule at the 5’ and 3’-termini, respectively **(Table S1)**, facilitating their chemical conjugation to protein binders harboring the non-natural amino acid azido-phenylalanine (AzF) via click chemistry. Oligo BL is hybridized to Left-T and initiates the synthesis of dsDNA by Bst2.0 polymerase. The resulting dsDNA is reacted with AzF-binders to form the Left-probe. Similarly, Oligo BR was hybridized to Right-T and then reacted with AzF-binders to form the Right-probe. A short spacer, poly-A_20_ (∼7 nm), is inserted in BL and BR between the DBCO and the annealing region to provide space for accommodating various target proteins.

**Figure 3.**
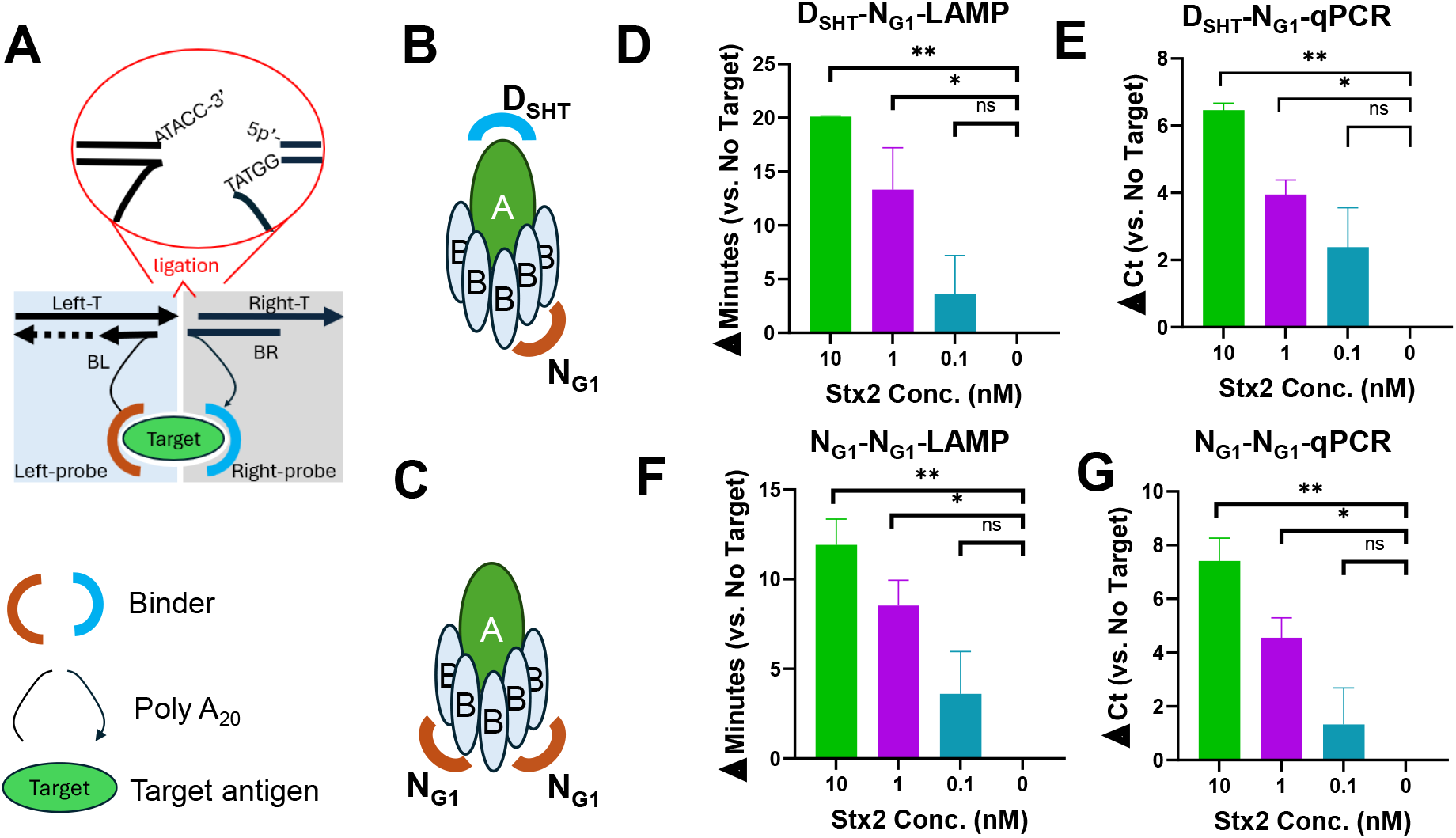
DISCO-LAMP.v1 detection of Stx2. **(A)** Schematic of DISCO-LAMP for the detection of target molecules via PLA. **(B, C)** Schematic of Stx2 detection using N_G1_-D_SHT_ pair or N_G1_-N_G1_ pair. **(D-G)** Calculated Δ Minutes or Δ Ct of Stx2 detection assay using LAMP or qPCR, respectively. The Δ Ct or Δ Minutes values are the average from two independent experiments carried out in technical duplicates. The limit of detection was determined using one-way ANOVA (n = 2 per group). Error bars indicate SEM and statistical significance was determined by Fisher’s LSD test versus No Target (*: p < 0.05; **: p < 0.005).

To demonstrate the utility of DISCO-LAMP for diagnostic application, we conjugated a pair of protein binders of Shiga toxin 2 (Stx2) to BL and BR to form the Left- and Right-probes. Shiga toxin-producing *E. coli* (STEC) is estimated to cause approximately 2.8 million acute illnesses annually^24^. The key virulence factor of STEC is Stx2, a typical AB_5_ family protein toxin comprised of an enzymatically active A-subunit and a non-toxic pentameric B-subunit^25, 26^. One of the Stx2 binder is D_SHT_, a designed ankyrin repeat protein (DARPin) previously engineered in our lab, which associates with the A-subunit of Stx2 and inhibits the toxin’s catalytic activity^27^. The other Stx2 binder is N_G1_, a nanobody specific for the B-subunit of Stx2^28^. N_G1_ and D_SHT_ were conjugated to BL/BR via click chemistry to form the Left-/Right-probe. Overall, 70-80% of AzF-N_G1_ reacted with Left-T/BL and ∼50% of AzF-D_SHT_/-N_G1_ reacted with Right-T/BR **(Figure S3)**. These probes were used without further purification to simplify the protocol and minimize reagent loss.

For PLA, the sensing solution contains Left- and Right-probes (10 nM of DNA each) and T4 DNA ligase (0.1 U/μL) in Buffer LP (1x T4 DNA ligase Buffer 1xPBS). Different concentrations of Stx2 were spiked in the sensing solution and the mixture (10 µL) was incubated at room temperature for 30 minutes to promote ligation and then inactivated at 70°C for 15 minutes. The samples were diluted 50-fold and 200-fold in ddH2O and 1 µL of the mixture were used in LAMP and qPCR (10 µL), respectively. The first pair comprises D_SHT_ and N_G1_, which target the A- and B-subunit, respectively **(Figure 3B)**. Considering the pentameric structure of the B-subunit, we also tested a second pair composed of N_G1_ conjugated to both Left- and Right-probes **(Figure 3C)**. Both pairs can reliably detect Stx2 at 1 nM via qPCR and LAMP, although detection at 10 nM is more statistically significant (p<0.005) (**Figure 3D-G**). The D_SHT_-N_G1_ pair seems to be slightly superior to the N_G1_-N_G1_ pair resulting in larger Δ Minutes values at all Stx2 concentrations. These results demonstrate proof-of-principle for protein detection by DISCO-LAMP/qPCR. However, neither pair can detect Stx2 at 0.1 nM, prompting us to further optimize the assay design to improve the detection sensitivity.

### DISCO-LAMP.v2 PLA detection of Stx2

DISCO-LAMP.v1 uses Bst2.0 polymerase to synthesize the dsDNA Left-Probe from oligo BL hybridized to oligo Left-T. Due to less than 100% efficiency of DNA hybridization and prematurely terminated DNA synthesis, partially singled stranded Oligo Left-T can potentially interfere with the subsequent primer annealing and LAMP/qPCR amplification, leading to high assay background and poor Stx2 detection sensitivity. To more efficiently produce the double-stranded Left-probe, we designed DISCO-LAMP.v2. In this new design, the F3 region in the original Left-T **(Figure 1B)** is replaced with F1c, which is complementary to the F1 region, resulting in a spontaneous hairpin structure in oligo Left-T.v2 **(Figure 4A)**. This hairpin structure is similar to that used by Pang, et al.^20^. This design negates the DNA synthesis step and should more efficiently produce the template for LAMP. To form Left-probe.v2, BL was first reacted with an AzF-binder **(Figure S4A)** and then hybridized with Oligo Left-T.v2 **(Table S1)**. A biotin molecule was added to the 3’ of Oligo Right-T, resulting in Right-T.v2. To create Right-probe.v2, BR was first reacted with AzF-binder before hybridizing with Right-T.v2.

**Figure 4.**
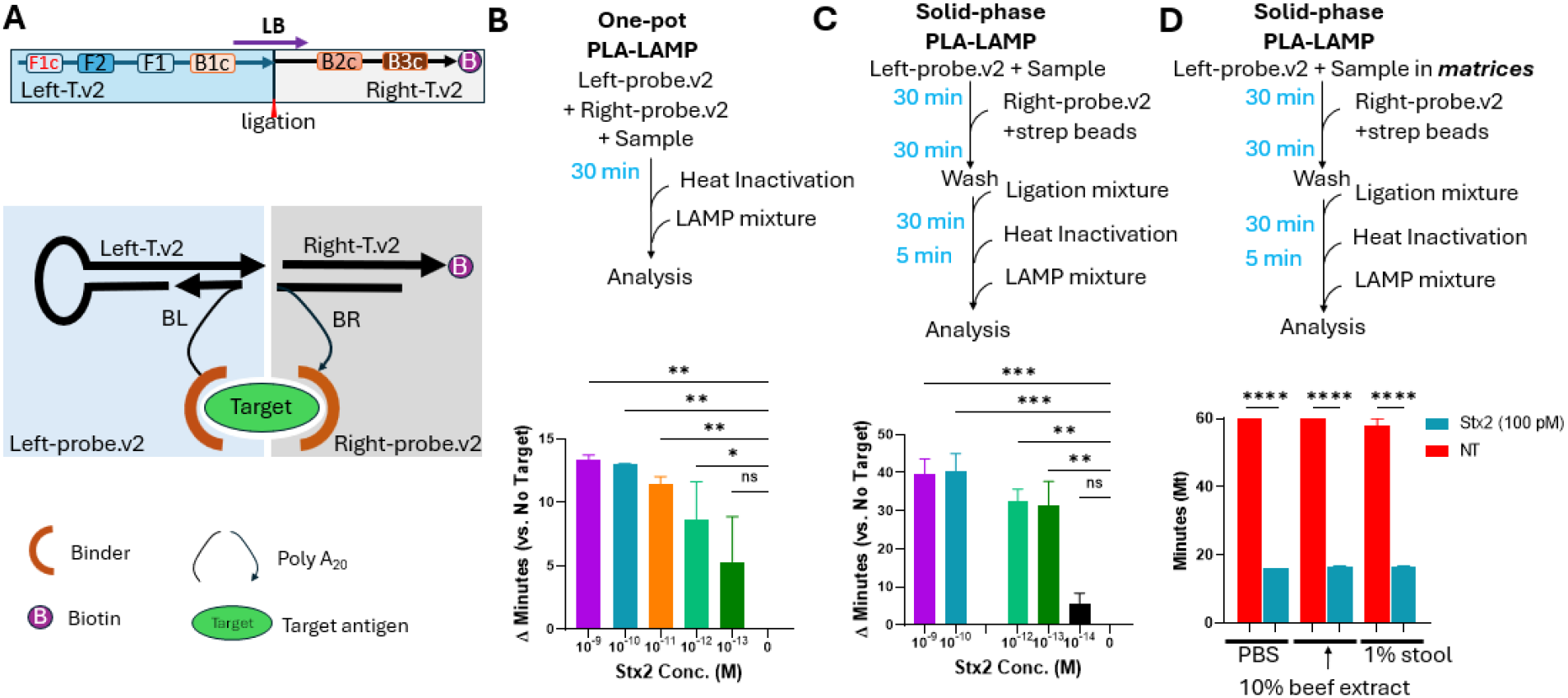
Detection of Stx2 using DISCO-LAMP.v2. **(A)** Schematics of DISCO-LAMP.v2 design and probe configuration. **(B)** One-pot homogenous detection of Stx2. Data represents the mean Δ Ct of biological replicates; error bars represent standard error of mean (SEM). **(C)** Solid-phase detection of Stx2 using bead-enriched ligation products, with data plotted as error bars representing the SEM of biological replicates. **(D)** Solid phase detection of Stx2 in different matrices, where Minutes (Mt) is determined by regression and error bars represent SEM. NT: No target with ligase control. For all panels, Δ Minutes and Mt values are the average from two independent experiments carried out in technical duplicates. Statistical significance was determined using one-way ANOVA (*n* = 2 per group) and Fisher’s LSD test relative to No Target (*: *p* < 0.05; **: *p* < 0.005, ***: *p* < 0.0005, ****: *p* < 0.0001).

Using the N_G1_-N_G1_ pair, we first developed a one-pot LAMP-PLA protocol for sensing Stx2. Left- and Right-probe.v2 (1 nM each) were mixed with T4 DNA ligase (0.1 U/µL) in Buffer LP spiked with different concentrations of Stx2 (10 µL total). After 30 minutes of incubation at room temperature and heat inactivation at 70°C for 15 minutes, LAMP mixture (90 µL) was added, and the mixture was incubated at 65 ºC for LAMP amplification **(Figure 4B)**. Using this one-pot format, DISCO-LAMP.v2 reliably detects Stx2 with LoD of 1 pM (∼90 pg/mL), which is ∼1000-fold better than DISCO-LAMP.v1 (∼1 nM). At concentrations between 1 pM and 100 fM, the detection of Stx2 is somewhat sporadic, likely due to the concentration of ligated DNA nearing the pseudo cut-off point for LAMP. The LoD of one-pot DISCO-LAMP.v2 is comparable to ELISA but takes < 2 hours and requires no washing step, pointing to a high translation potential.

Using the same N_G1_-N_G1_ pair and taking advantage of the biotin molecule on oligo Right-T.v2, we also developed a solid-phase sensing protocol with further improved Stx2-detection sensitivity. In this protocol, Left-probe.v2 (1 nM) was incubated in buffer PTBD (1x PBS supplemented 0.05% Tween 20, 0.2% BSA and 0.3 mg/mL salmon sperm DNA) in the absence or presence of Stx2 at room temperature for 30 minutes before the addition of Right-Probe.v2-functionalized streptavidin magnetic beads to pull-down Stx2 and associated Left-probe.v2. After extensive washing with PTBD, the beads were resuspended in Buffer LP supplemented with T4 DNA ligase (0.2 U/µL) to initiate the PLA reaction. After 30 minutes of incubation at room temperature, the beads were washed and heat inactivated, and the supernatant was used for LAMP analysis. This solid-phase LAMP protocol achieved reliable detection of Stx2 with LoD of 100 fM (∼9 pg/mL), making it 10 times more sensitive than the one-pot LAMP protocol and 10,000-fold more sensitive than DISCO-LAMP.v1) **(Figure 4C)**.

Although more time consuming and complex, a major advantage of the solid-phase sensing protocol is the separation of the DNA/protein assembly step from the ligation step, which is critical as the activity of the ligase is often compromised in challenging matrices. To evaluate the performance of our solid-phase PLA-LAMP protocol in complex matrices, we repeated the assay and incubated the Left-probe.v2 in PTBD spiked with 10% beef extract or 1% stool extract in the absence or presence of Stx2 (100 pM, ∼9 ng/mL). Stool and beef were chosen as representative matrices because they are frequently contaminated with STEC. Similar LAMP amplification efficiency was observed for samples incubated in PBS or 10% beef/ 1% stool extract **(Figure 4D)**, demonstrating the compatibility of DISCO-LAMP.v2 and the solid-phase sensing protocol with 10% beef and 1% stool matrices. Higher concentrations of beef/stool extract obfuscate Stx2 sensing, likely due to the presence of nuclease in these matrices.

### DISCO-LAMP.v2 PLA detection of SARS-CoV-2

To demonstrate the utility of our DISCO-LAMP in another clinically relevant target, we evaluated its ability to detect SARS-CoV-2 spike protein. We previously engineered a panel of DARPins able to potently neutralize SARS-CoV-2^29^. From the same screen, DARPin S16 (D_S16_) emerged as a strong binder targeting a highly conserved epitope on Spike protein. Although D_S16_ was engineered using exclusively Wuhan-1 Spike, which differs significantly from later viral variants, in ELISA, D_S16_ can efficiently bind spike protein from diverse SARS-CoV-2 strains, including Wuhan-1 and Omicron, with estimated EC_50_ < 1 nM in ELISA **(Figure 5A)**. Additionally, by BLI, the K_a_ of D_S16_ to full-length Spike of Wuhan-1 and Omicron strains are 3.7×10^4^ (1/Ms) and 3.3×10^4^ (1/Ms), respectively. The K_D_ values are below the detection limit, likely due to the avidity effect since the Spike protein is a trimer **(Figure 5B, C)**. Since the spike protein is a trimer, D_S16_ was used to form both Left- and Right-probes in DISCO-LAMP.v2 **(Figure 5D)**. The conjugation efficiency of both probes is estimated to be ∼50% (**Figure S4B**). Using the one-pot sensing protocol **(Figure 4B)**, DISCO-LAMP.v2 efficiently detected Wuhan-1 and Omicron spike protein at 1 nM **(Figure 5E, F)**. Lower concentrations of Spike protein were not tested. These data demonstrate the potential of DISCO-LAMP.v2 for sensing other clinically relevant targets.

**Figure 5.**
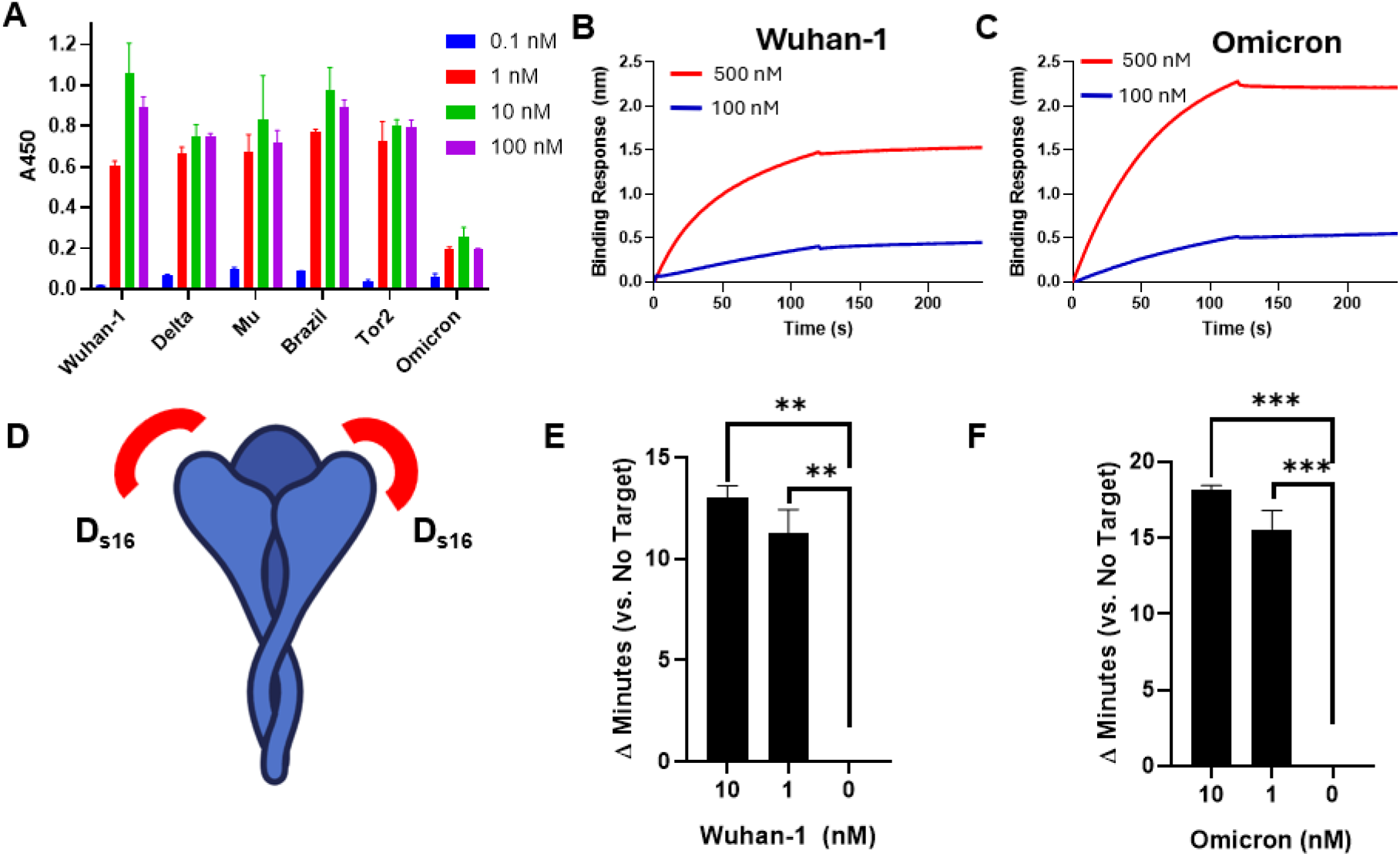
DISCO-LAMP.v2 detection of SARS-CoV-2 Spike protein. **(A)** Relative binding of Wuhan-1 and Omicron Spike Proteins by D_S16_, determined using ELISA. BLI sensograms of D_S16_ to Spike proteins from Wuhan-1 **(B)** and Omicron **(C). (D)** Schematic of Spike detection by D_S16_. **(E, F)** Calculated Δ Minutes of DISCO-LAMP of Wuhan-1 and Omicron spike detection versus no target. The Δ Minutes values are the average from two independent experiments carried out in technical duplicates. The limit of detection was determined using one-way ANOVA (n = 2 per group). Error bars indicate SEM, and statistical significance was determined by Fisher’s LSD test versus No Target (*: p < 0.05; **: p < 0.005, ***: p < 0.0005).

## Discussion

PLA is an effective method for detecting proteins and other antigens with high sensitivity and specificity. LAMP serves as an ideal detection tool since it enables quick, exponential DNA amplification at a constant temperature, making it especially useful for field applications and point-of-care testing. In the PLA–LAMP context, it is essential that the probe halves do not lead to LAMP amplification to minimize the background signal. The exponential nature of LAMP and the use of multiple primers make it challenging to design a discontinuous LAMP system as rare off-target priming events can be rapidly amplified to give rise to substantial background signals. Our initial efforts focused on splitting the LAMP template in the middle, separating F1–F3 from B1–B3. However, even after extensive primer/template redesign, we could not sufficiently suppress the background signal from single halves. Finally, we landed a new design that split the template at a loop primer (LB) annealing region and developed DISCO-LAMP **(Figure 1)**. In this configuration, individual halves at up to 10 nM concentration fail to produce detectable LAMP signal after 60 minutes **(Figure S1)** while the full-length (ligated) template can be at 1 fM. Although this detection sensitivity is slightly weaker than that of the most highly optimized LAMP or qPCR assays, it is sufficiently sensitive for many diagnostic applications.

Proof-of-concept experiments using a pair of complementary oligonucleotides to mimic target induced proximity demonstrated that DISCO-LAMP.v1 can be efficiently triggered by duplex formation and read out by both qPCR and LAMP. Systematic optimization showed that the ligation time and ligase concentration are relatively forgiving, whereas buffer composition—particularly the inclusion of 1xPBS—plays a major role in improving the signal-to-noise ratio **(Figure 2)**. The platform was then applied to protein detection using Shiga toxin 2 (Stx2), a key virulence factor of STEC. Two previously engineered protein binders (D_SHT_ and N_G1_) were conjugated to the DNA probe via copper free click chemistry. These experiments established the feasibility of wash free protein detection but exhibited limited sensitivity (LoD 1 nM, **Figure 3**), likely stemming from a modest signal to noise ratio driven by partially single-stranded LAMP template DNA.

To address these limitations, we refined the template design in DISCO-LAMP.v2 so that one probe arm carries a preformed LAMP hairpin structure and the other incorporates a biotin moiety **(Figure 4)**. The new hairpin design significantly improved the signal-to-noise ratio, and the biotinylated arm enables streptavidin mediated immobilization and washing steps, which further reduce background by physically removing free probes. Under otherwise similar reaction conditions, DISCO-LAMP.v2 achieved 10^3^-10^4^-fold improved Stx2 detection sensitivity with LoD of 1 pM–100 fM. The same architecture also enabled the detection of SARS-CoV-2 spike protein. These gains highlight the combined importance of template design and physical purification in achieving high sensitivity target detection.

A critical dictator of PLA efficiency is the target-binding included proximity of the DNA probes, specifically the ligation partners (3’ of the Left-T (sticky end) and the 5’ of Right-T). Due to the limited repertoires of aptamer-based binders, protein-based binders, mainly antibodies, have been extensively used in PLA assays. Unlike aptamers, which can be chemically synthesized as a natural extension of the DNA probe, special techniques are needed to conjugate a protein binder to the DNA probe. Since both protein and DNA can be easily biotinylated, streptavidin has been extensively used as the bridge to connect a protein binder to a DNA probe^30-32^. The very high affinity between biotin and streptavidin enables the formation of a stable protein-streptavidin-DNA probe molecule. However, the tetrameric structure of streptavidin inevitably increases the length and size of the linker, thus reducing the local DNA probe concentration and potentially reducing the ligation efficiency. In addition, since each streptavidin contains four biotin binding sites, it is impossible to obtain a homogenous population of protein-DNA conjugates. In the past decades, alternative methods have been developed to create protein-DNA conjugates without introducing bulky intermediates, including strain promoted azide alkyne cycloaddition (SPAAC)^33-36^, inverse electron demand Diels alder (IEDDA) reactions^37^, intein ligation^38^ and maleimide-thiol reactions^11, 12, 14, 39-43^. Among these, copper-free click chemistry between DBCO and azide (N_3_) has been steadily gaining popularity due to its high efficiency, stability and specificity^44^, and is adopted in our DISCO-LAMP. When both binders are associated with the same target molecule, the maximum molecular space occupied by ligation partners is determined by the length of the spacer in BL and BR (*e*.*g*. poly-A_20_ (∼7 nm) and the size of the binder molecules. This direct and site-specific click chemistry-mediated protein–DNA conjugation method allows precise control of linker length and composition, making it feasible to customize linker lengths for different targets to achieve maximum PLA efficiency. This modular, noncanonical amino acid– based strategy provides an effective method to produce protein-DNA probes.

The current sensitivity of DISCO-LAMP.v2 for Stx2 (LoD ∼100 fM, ∼9 pg /mL) slightly surpasses that of established ELISA formats (LoD 25-50 pg/mL^45^). In addition, our method takes < 2 hours and provides a simpler and more user-friendly workflow. The platform also performs robustly in 10% beef and 1% stool matrices that are frequently contaminated with STEC, pointing to its utility in real world sample types. Taken together, this work establishes: (i) a discontinuous LAMP template and primer design intrinsically compatible with PLA and (ii) a clear design in which a preformed hairpin and carefully tuned linker lengths jointly control local concentration and assay sensitivity. The detection of Stx2 and SARS-CoV-2 spike protein demonstrates the versatility of DISCO-LAMP. Future efforts will focus on fine tuning the linker length and sampling alternative protein binders with better target-binding affinity and specificity to further improve LoD for diverse diagnostic applications.

## Summary

In conclusion, this work establishes a discontinuous DISCO - LAMP template and primer architecture that is compatible with PLA, enabling robust amplification of ligated templates while minimizing background from individual probe halves. Using this framework, DISCO-LAMP sensing protocols achieve femtomolar-range limits of detection for Stx2, with performance comparable to ELISA yet in a faster, isothermal and potentially field-deployable format. The study also introduces D_S16_, a newly engineered DARPin that targets a conserved SARS-CoV-2 spike epitope and supports spike detection. We believe that DISCO-LAMP can potentially be extended to other diagnostic applications.

## Materials and Methods

### Protein expression and purification

All proteins were expressed in *E. coli* BL21(DE3) cells except for AzF-N_G1_, which is a nanobody and was expressed in Shuffle® T7 cells (NEB, Cat# C3029J). For expression of AzF-D_SHT_, AzF-D_S16_ and AzF-N_G1_, the *E. coli* were transformed with pEVOL-AzFRS^46^ and the appropriate expression plasmid. The cells were cultured in Luria-Bertani (LB) broth supplemented with 12.5 μg/mL chloramphenicol and 50 μg/mL kanamycin. Once the culture reached OD_600_ ∼0.5, 0.2 % ( w/v) L-(+)-arabinose (Chem-Impex Int’l. Inc., Cat#01654) was added, and the culture was transferred to an 18°C shaking incubator. Thirty minutes later, isopropyl b-d-1-thiogalactopyranoside (IPTG, final 0.5-1 mM, Goldbio, Cat# I2481C) and 4-Azido-L-phenylalanine (Chem-Impex Cat# 06162), final 1 mM were added to induce target protein expression. For expression of D_S16_, the culture was induced with 0.1 mM IPTG at OD_600_ ∼0.7 and grown at 18ºC. The cells were harvested the next day, and the 6xHis-tagged target proteins were purified using immobilized metal affinity chromatography (IMAC) with gravity Ni-NTA beads (PureCube 100 Ni-NTA Agarose, Cat# 74705) following standard protocol. The eluted protein was buffer exchanged into 1x PBS (Fisher Cat# BP399) and concentrated via ultrafiltration (Millipore, MWCO 10 kDa or 3k Da, Cat# UFC9010 or Cat# UFC9003). The purity of each protein was analyzed using 12% Mini-PROTEAN® TGX™ Precast Protein Gels (Bio-Rad, Cat# 4561043) and are all > 90% **(Figure S3)**. The protein concentration was quantified using Pierce BCA Protein assay (Thermo Scientific, Cat# PI23227). Stx2 was expressed and purified as previously described^27^.

### Preparation of full-length DNA template

Full-length DNA template was prepared by ligation of oligos Left-T and Right-T with a splint oligo, splint-LAMP **(Table S1)**. The ligation product was PCR amplified using primers qF and qR2. The PCR product was gel purified, quantified using NanoDrop, and stored in aliquots at -20ºC.

### Preparation of Left-/Right-probes

Initial DISCO-LAMP probes **(Figure 2A)**: To prepare the Left-probe, oligo D1 (2 μM) was first hybridized to Left-T (2 µM) in 1x isothermal LAMP buffer (NEB) using the following hybridization program: 95°C for 1 minute, 70°C for 5 minutes, ramp at -0.1°C/s to room temperature with 5 minutes hold every 5°C when the temperature reaches below 50°C. Bst 2.0 (NEB, Cat # M0538S) and dNTPs (final 0.4 mM, NEB) were then added and the mixture was incubated at 65°C for 60 minutes. To prepare the Right-probe, oligo D2 or ND2 (1 µM) was hybridized to Right-T (1 µM) in PBS using the same program. The resulting hybridized DNA probes were aliquoted and stored at -20°C until use.

DISCO-LAMP.v1 **(Figure 3A)**: To prepare the Left-probe, oligo BL (40 μM) was first hybridized to Left-T (40 µM) in 1x isothermal LAMP buffer (NEB) with 1mM of dNTPs using the same hybridization program Bst 2.0 was then added and the mixture was incubated at 65°C for 60 minutes. The resulting double-stranded DNA was reacted with AzF-D_S16_ (final 10 µM) or AzF-N_G1_ (final 15 µM) at 2:1 molar ratio in PBS at room temperature overnight to form the appropriate Left-probe and stored at -20°C in 50% glycerol until use. To prepare the Right-probe, oligo BR (45 µM) was hybridized to Right-T (45 µM) in PBS using the same hybridization program. The resulting hybridized DNA was reacted with AzF-D_S16_ (final 20 μM), AzF-D_SHT_ (final 30 μM), or AzF-N_G1_ (final 10 μM) at 1:1 molar ratio in PBS at room temperature overnight and stored at -20°C in 50% glycerol until use.

DISCO-LAMP.v2 **(Figure 4A)**: To prepare the Left-probe.v2, oligo BL (∼70 μM) was first reacted with AzF-D_S16_ or AzF-N_G1_ at 1:1 molar ratio in PBS at room temperature overnight, and then diluted and hybridized to oligo Left.T.v2 (final 2 µM each). For Right-probe.v2, oligo BR (∼70 μM) was reacted with AzF-D_S16_ or AzF-N_G1_ at 1:1 molar ratio in PBS at room temperature overnight, and then hybridized to N_G1_-BR or D_S16_-BR. Aliquots were stored at -20°C in 50% glycerol until use.

All probes were analyzed via SDS-PAGE gels with SybrGreen staining to visualize DNA conjugation and Coomassie blue staining for protein detection. Reaction efficiency was estimated using ImageJ gel analysis.

### Target detection

For the initial DISCO-LAMP proof of concept, the PLA was carried out in Buffer L (1x T4 DNA ligase Buffer, 100 pM Left-probe (D1), and 10 nM of Right-probe (D2 or ND2) in the absence or presence of 1x PBS and various concentrations of T4 DNA ligase. The mixtures were incubated at room temperature for 1-30 minutes, heat inactivated at 70°C for 15 minutes, and analyzed using qPCR and LAMP. For LAMP detection, 1 μL of the reaction was transferred to 9 μL LAMP mixture (0.2 µM of F3, 0.2 µM of B3, 1.6 µM of FIP, 1.6 µM of BIP, 0.4 µM of LB, 0.4 mM of dNTPs, 3.2 U of Bst Warmstart® 2.0 DNA polymerase, 0.2X EvaGreen dye (Biotium, Cat #31019), 4mM MgSO_4_ (NEB), and 1×isothermal amplification buffer (NEB)). The reaction was carried at 65°C in a qPCR instrument (Bio-Rad CFX Duet Real-Time PCR). For qPCR detection, 1 μL of the reaction was transferred to 9 μL qPCR mixture (200nM qF and qR in 1x Forget-Me-Not™ EvaGreen® qPCR Master Mix (Cat #31041)). The mixture was analyzed in a qPCR instrument using the following program (95°C for 2 min, followed by 40 cycles of 95°C for 15 s, 60°C for 30 s and 72°C for 30 seconds with a final extension of 72°C for 1 minute).

For DISCO-LAMP.v1, the PLA was carried out in Buffer LP (1xPBS and 1x T4 DNA ligase Buffer (NxGen)) supplemented with 10 nM of Left-/Right-probe, 0.1 U/μL of T4 DNA ligase (NxGen) and different concentration of Stx2. The mixture was incubated at room temperature for 30 minutes and inactivated at 70°C for 15 minutes. For LAMP detection, the mixture was diluted 50-fold using ddH2O and 1 μL of the diluted mixture was transferred to 9 μL LAMP mixture. For qPCR detection, the ligation mixture was diluted 200-fold in ddH2O and 1 μL of the diluted mixture was transferred to 9 μL qPCR mixture.

For DISCO-LAMP.v2 detection of Stx2 in one-pot format, the PLA was carried in the same Buffer LP except that 1 nM of Left-/Right-probe.v2 was used. For Omicron detection, the probe concentration was 2.5 nM while the Wuhan-1 probe concentration was maintained at 1 nM. After inactivation, 9-fold reaction volume of LAMP mixture (excluding F3) was added, and reaction was carried out at 65°C in a qPCR instrument.

For DISCO-LAMP.v2 in the solid-phase, different concentrations of Stx2 were incubated with N_G1_-functionalized Left-probe.v2 (1 nM) in buffer PTBD (1x PBS supplemented with 0.05% tween-20, 0.2% BSA, and 0.3 mg/mL salmon sperm DNA) at room temperature for 30 minutes. The mixture was then incubated with N_G1_-Right-probe.v2 -functionalized MyOne streptavidin-coated magnetic beads (Thermo Fisher Scientific Cat# 65601) at room temperature for 30 minutes. The beads were thoroughly washed in PTBD and finally incubated in Buffer LP supplemented with 0.2 U/μL of T4 DNA ligase at room temperature for 30 minutes and inactivated at 95°C for 5 minutes. The supernatant was removed, and the beads were resuspended in 1xNEB ThermoPol buffer (Cat# B9004S) and the amount of ligated DNA in the supernatant was subsequently amplified using LAMP.

For solid-phase PLA-LAMP detection of Stx2 in complex matrices, the PTBD was spiked with 1% stool (w/v) or 10% beef extract (v/v). Beef extract was obtained by collecting liquid released by compression of beef purchased from a local retailer. Stool was obtained from Innovative Research (SKU: IRHUSFM1). Stool was resuspended in 1xPBS to a final concentration of 10%, centrifuged at 12,000g for 5 min, and the resulting supernatant was aliquoted and stored at -20 °C.

All Ct and Minutes values were determined using the Cfx-threshold method from CFX Maestro Version 2.3 software unless stated otherwise.

### Characterization of D_S16_

D_S16_ was derived from phage panning against Wuhan-1 spike protein^29^ and harbors a Myc-tag at the N-terminus. The ability to bind spike proteins was evaluated using ELISA. Briefly, high binding ELISA (Nunc MaxiSorp, Fisher Scientific, Cat# 50-712-278) plates were coated with various Spikes proteins (4 µg/mL) overnight at 4ºC. The next day, the plate was blocked using PBS supplemented with 2% BSA and 0.1% Tween-20 before the addition of serially diluted D_s16_, and the amount of immobilized D_S16_ after washing was detected using chicken anti-Myc HRP antibody (Immunology Consultants Laboratory, Cat# cmyc-45P-Z, 1: 20,000 dilution from stock) and TMB substrate. Absorbances at A450 were recorded using Tecan Infinite M200 PRO microplate reader.

To measure the binding affinity of D_S16_ to spike protein, purified AzF-D_S16_ (final 2 mg/mL) was first reacted with biotin-PEG4-DBCO (Click-Chemistry Tools Cat # A105-5) at 1:10 molar ratio in 1xPBS and incubated at 4 ºC for 24-48 hours with rotation. Excess DBCO-biotin was removed using Zeba spin desalting columns (7K, MWCO, ThermoFisher) twice. The resulting biotin-D_S16_ (250nM) was resuspend in PBS with 0.1% BSA, loaded on Streptavidin sensors (Sartorius #18-5019) on a Fortebio BLItz instrument (Sartorius, Goettingen, Germany) and analyzed using 100 nM or 500 nM Wuhan-1 or Omicron Spike protein in the same buffer. Every measurement consisted of 30 s of equilibration, 120 s of D_S16_ loading, 30 s of baseline, 120 s of Spike association, and 120 s of Spike dissociation. BLItz Pro 1.3 software was used for binding detection and analysis. Due to the avidity effect, no Spike dissociation was observed.

## Author Contributions

Z.C., B.T., and R.S. conceived the project; B.T., R.S., V. C., W.C., K.Y., E.W., J.H., A.F. and C.M. conducted the experiments; B.T. and Z.C. wrote the manuscript; All authors have read and agreed to the published version of the manuscript.

## Acknowledgements

The following reagents were obtained through BEI Resources, NIAID, NIH: Spike Glycoprotein (Stabilized) from SARS Coronavirus, **Tor2** with C-Terminal Histidine and Strep® II Tags, Recombinant from HEK293 Cells, NR-53590; Spike Glycoprotein (Stabilized) from SARS-Related Coronavirus 2, **Delta** Variant with C-Terminal Histidine and Avi Tags, Recombinant from HEK293 Cells, NR-55614; Spike Glycoprotein (Stabilized) from SARS-Related Coronavirus 2, B.1.621 Lineage (**Mu** Variant) with C-Terminal Histidine and Avi Tags, Recombinant from HEK293 Cells, NR-55712; Spike Glycoprotein (Stabilized) from SARS-Related Coronavirus 2, P.1 Lineage (**Brazil**) with C-Terminal Histidine and Avi Tags, Recombinant from HEK293 Cells, NR-55307; Spike Glycoprotein (Stabilized) from SARS-Related Coronavirus 2, BA.2 Lineage (**Omicron** Variant) with C-Terminal Histidine and Avi Tags, Recombinant from HEK293 Cells, NR-56517; and Spike Glycoprotein (Stabilized) from SARS-Related Coronavirus 2, **Wuhan-Hu-1** with C-Terminal Histidine and Twin-Strep® Tags, Recombinant from HEK293 Cells, NR-52724.

We would like to express gratitude to members of the Chen research team for healthy discussions and constructive feedback.

## Notes

### Competing Interest Statement

The authors have declared no competing interest.

### Funding

- NIH: GM138811, DP2AI136600, R01Al180650
- Texas A&M TPT

