## Supplemental Information for "DISCO-LAMP: A Novel discontinuous LAMP assay for isothermal antigen detection"

**Supplementary Information**

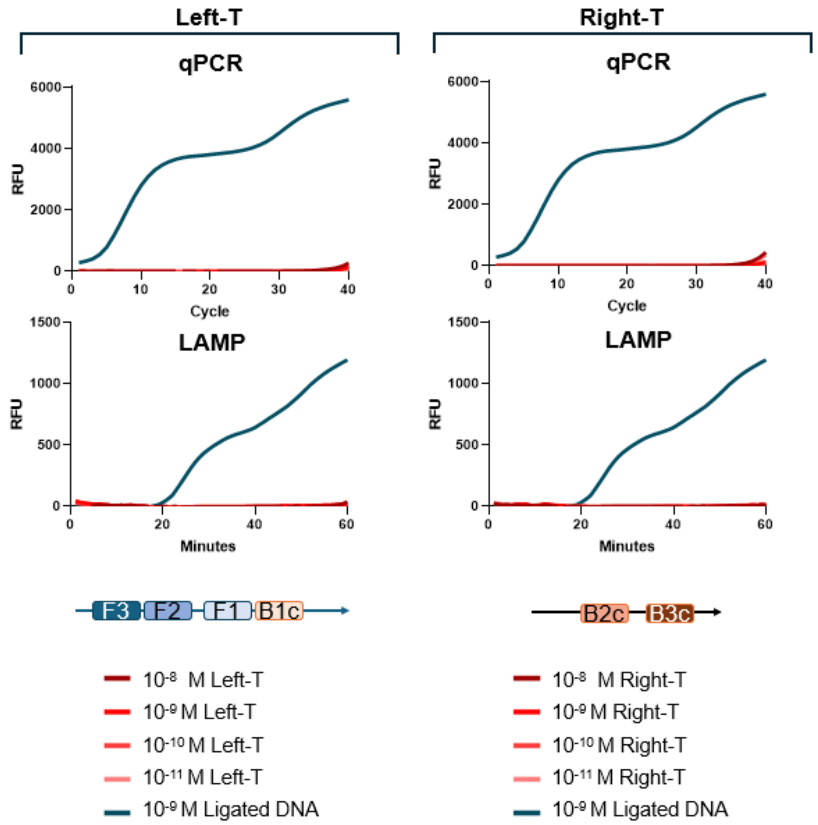


**Figure S1. Comparison of amplification efficiency of full-length template and individual Left-/Right-T oligos by qPCR and LAMP.** Representative amplification curves comparing full-length (ligated DNA) template and individual Left-/Right-T oligos amplified by qPCR and LAMP. Representative data from at least 2 experiments were shown.

**A**

| Name | Sequence |
|---|---|
| D1 | GGTTCCTTAATACGTC AAAAA GTACCTACCCCCGTCCTTTTG |
| D2-6 | ACCCCACCAACTAGCTGATATGGCTGA**GGTATA**GAAAAAGACGTATTAAGGAACC |
| ND2-6 | ACCCCACCAACTAGCTGATATGGCTGA**GGTATA**G AAAAA |
| D2-5 | ACCCCACCAACTAGCTGATATGGCTGA**GGTAT**TG AAAAAGACGTATTAAGGAACC |
| ND2-5 | ACCCCACCAACTAGCTGATATGGCTGA**GGTAT**TG AAAAA |
| D2-4 | ACCCCACCAACTAGCTGATATGGCTGA**GGTA**ATGAAAAA GACGTATTAAGGAACC |
| ND2-4 | ACCCCACCAACTAGCTGATATGGCTGA**GGTA**ATG AAAAA |
| D2-0 | ACCCCACCAACTAGCTGATATGGCTGACCATATGAAAAA GACGTATTAAGGAACC |
| ND2-0 | ACCCCACCAACTAGCTGATATGGCTGACCATATG AAAAA |

Regions annealing to the sticky end at the 3' of Left-T are shown in red.

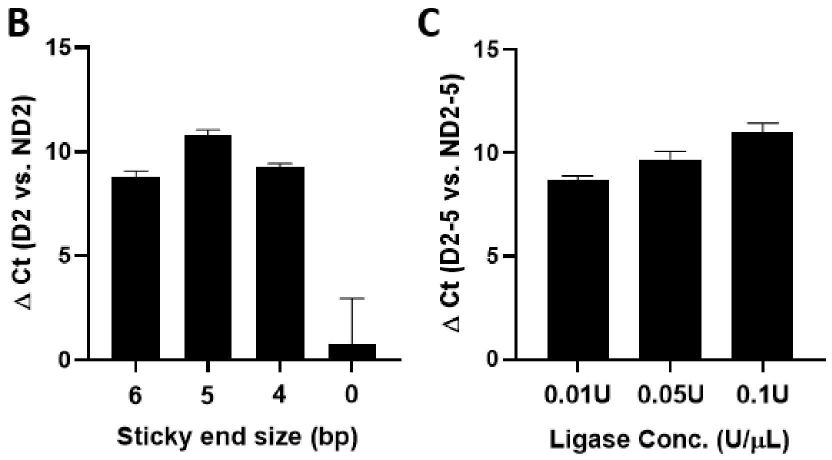


**Figure S2. DISCO-LAMP assay optimization. (A)** Oligo sequences. Red letters indicate the regions forming the sticky end with Left-T. The impact of the length of sticky end **(B)** and T4 DNA ligase concentration **(C)** on the PLA efficiency determined by qPCR.

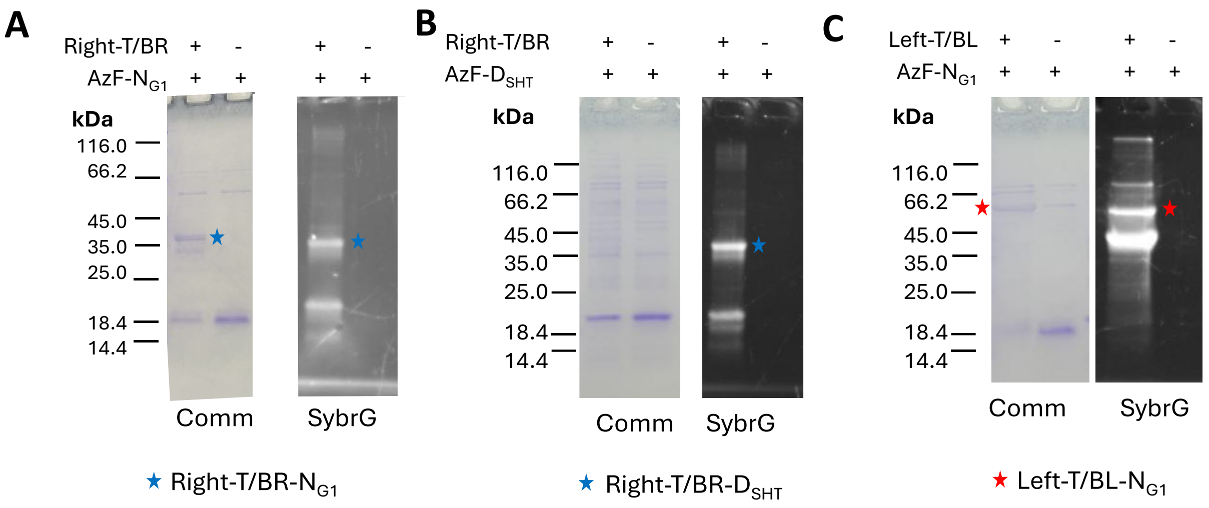


**Figure S3. Characterization of Left-/Right-probes.** Reactions of Right-T/BR with AzF-$N_{G1}$ **(A)**, Right-T/BR with AzF-$D_{SHT}$ **(B)** and Left-T/BL with AzF-$N_{G1}$ **(C)** analyzed on 12% SDS-PAGE gels. The gels were first stained for DNA using SybrGreen (SybrG), imaged, and then stained for protein using Coomassie blue (Comm). Equal amounts of proteins were loaded in the adjacent lanes, and the reaction efficiency is estimated from the reduction of the band intensity of unreacted protein.

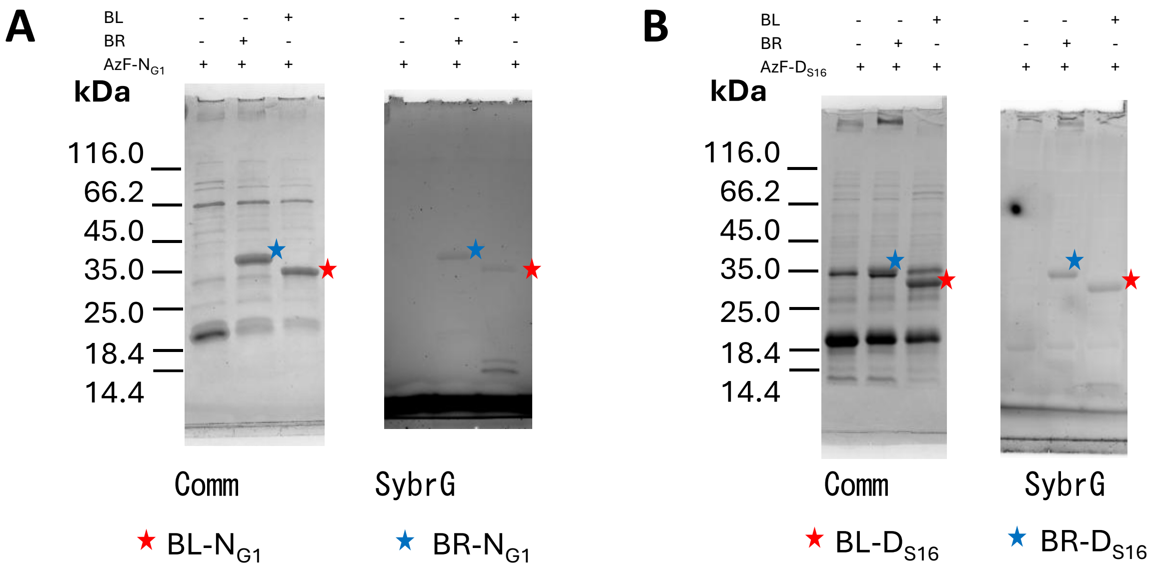


**Figure S4. Characterization of oligo BL/BR conjugation to $N_{G1}$/$D_{S16}$.** The gels were first stained for DNA using SybrGreen (SybrG), imaged, and later strained for protein using Coomassie blue (Comm). Equal amounts of proteins were loaded in each 12% SDS-PAGE gel, and the conjugation efficiency is estimated from the reduction of the band intensity of unreacted protein.

**Table S1. Primer sequences.**

| Lab Name | Sequence |
|---|---|
| Left-T.v2 | TGCACAACATGGGGGATCATGTGCATCGTGGTGTCACAGC TCCGGTTCCCAACGATCAAGGCGAGTTACATGATCCCCCA TGTTGTGCAAAAAAGCGGTTAGCAAAAGGACGGGGGTAGG TACAAGTATACC |
| Right-T.v2 | Phos/TCAGCCATATCAGCTAGTTGGTGGGGTATGGTTGTCA GAAGTAAGTTGGCGGC-AAAAA-biotin-teg |
| BL | /5DBCOTEG/AAAAAAAAAAAAAAAAAAAAGTACCTACCC CCGTCCTTTTG |
| BR | ACCCCACCAACTAGCTGATATGGCTGAGGTATTGAAAAAA AAAAAAAAAAAAAA /3DBCON/ |
| qF | CAACGTTGTTGCCATTGCTA |
| qR | ACCCCACCAACTAGCTGA |
| splint-LAMP | AGCTGATATGGCTGA GGTATACTTGTACCT |
| qR2 | CCAACTTACTTCTGACAACCAT |

**Table S2. Protein sequences.**

| Name | Target | Sequences |
|---|---|---|
| AzF-SHT | Stx2-A subunit | MG*EQKLISEEDLGSDLGKKLLEAARAGQDDEVRILVANGADVNAGDPFGF TPLHLAALYGHLEIVEVLLKNGADVNAHEEYGFTPLHLAAVVSHMEIVEVLLN NGADVNARDNQGGSPLHLASHTGHLEIVEVLLKQGADVNAQDKFGKTAYD ISIDIGNEDLAEILQSSSKLAAALE**HHHHHH** |
| AzF-G1 | Stx2-B subunit | MGSS*GSSQVQLVESGGGLVQPGESLRLSCVASASTFSTSLMGWVRQAPG KGLESVAEVRTTGGTFYAKSVAGRFTISRDNAKNTLYLQMNSLKAEDTGVYYC TAGAGPIATRYRGQGTQVTVSSAHHSEDPSSSSLE**HHHHHH** |

*** Amber codon for non-canonical amino acid incorporation**
**Myc tag**
